# Prediction of antidepressant treatment response with thalamo-somatomotor functional connectivity revealed by generalizable stratification of depressed patients

**DOI:** 10.1101/2024.05.11.593664

**Authors:** Yuto Kashiwagi, Tomoki Tokuda, Yuji Takahara, Yuki Sakai, Junichiro Yoshimoto, Ayumu Yamashita, Toshinori Yoshioka, Koichi Ogawa, Go Okada, Yasumasa Okamoto, Mitsuo Kawato, Okito Yamashita

## Abstract

Major depressive disorder (MDD) is diagnosed based on symptoms and signs without relying on physical, biological, or cognitive tests. MDD patients exhibit a wide range of complex symptoms, and it is assumed that there are diverse underlying neurobiological backgrounds, possibly composed of several subtypes with relatively homogeneous biological features. Initiatives, including the Research Domain Criteria, emphasize the importance of biologically stratifying MDD patients into homogeneous subtypes using a data-driven approach while utilizing genetic, neuroscience, and cognitive information. If biomarkers can stratify MDD patients into biologically homogeneous subtypes at the first episode of depression, personalized precision medicine may be within our scope. Some pioneering studies have used resting-state functional brain connectivity (rs-FC) for stratification and predicted differential responses to various treatments for different subtypes. However, to our knowledge, little research has demonstrated reproducibility (i.e., generalizability) of stratification markers in independent validation cohorts. This issue may be due to inherent measurement and sampling biases in multi-site fMRI data, or overfitting of machine learning algorithms to discovery cohorts with small sample sizes, i.e., a lack of appropriate machine learning algorithms for generalizable stratification. To address this problem, we have constructed a multi-site, multi-disorder fMRI database with prospectively and retrospectively harmonized data from thousands of samples and proposed a hierarchical supervised/unsupervised learning strategy. In line with this strategy, our previous research first developed generalizable MDD diagnostic biomarkers using this fMRI database of MDD patients via supervised learning. The MDD diagnostic biomarker determines the importance of thousands to tens of thousands of rs-FCs across the whole brain for MDD diagnosis. In this study, we constructed stratification markers for MDD patients using unsupervised learning (Multiple co-clustering) with a subset of top-ranked rs-FCs in the MDD diagnostic biomarker.

We developed a method to evaluate the clustering stability between two independent datasets as a generalization metric of stratification biomarkers. To discover stratification biomarkers with high stability across datasets, we utilized two multi-site datasets with substantial differences in data acquisition facilities and fMRI measurement protocols (Dataset-1: a dataset of 138 depressed patients obtained with a unified measurement protocol across three facilities; Dataset-2: a dataset of 181 depressed patients obtained with non-unified measurement protocols across four facilities, distinct from Dataset-1). Starting from several diagnostic biomarkers, we constructed some stratification markers and identified the stratification biomarker with the highest clustering stability between the two datasets. This stratification biomarker was based on several rs-FCs between the thalamus and the postcentral gyrus, and the MDD subgroups stratified by this biomarker showed significantly different treatment responsiveness to a selective serotonin reuptake inhibitor (SSRI).

By narrowing down whole-brain rs-FCs using MDD diagnostic biomarkers and further dividing the rs-FCs using multiple co-clustering, the feature dimension was significantly reduced, thereby avoiding overfitting to the training data and successfully constructing stratification biomarkers that are highly stable between independent datasets, i.e., have generalizability. Furthermore, the correlation between MDD subgroups and antidepressant treatment response was demonstrated, suggesting the potential for achieving personalized precision medicine for MDD.

## INTRODUCTION

Diagnosis in psychiatric disorders including major depressive disorder (MDD), in contrast to most medicine, remains restricted to subjective symptoms and observable signs without resorting to physical or biological examination (1). Although MDD is often diagnosed clinically as a single disorder, it is generally assumed that underlying neurobiological backgrounds are diverse, as MDD patients exhibit a wide variety of combined symptoms (2). Because treatment modalities may not be tailored to the pathophysiological factors of individual patients, treatment response is also highly variable (3), and it has been reported to take 2-4 years for approximately 60% of MDD patients to achieve remission (35, 36, 37). If objective biomarkers could stratify MDD patients into neurobiologically homogeneous subtypes, it might open up the possibility of personalized precision medicine (4). In other words, by selecting the optimal treatment for each subtype identified by stratification biomarkers at the initial depressive episode, it may be possible to shorten the time required for remission.

The National Institute of Mental Health (NIMH) launched the Research Domain Criteria (RDoC) project in 2009 to ultimately provide a framework for classification based on dimensions of neurobiology and observable behavior (5). RDoC classification conceptualizes mental illnesses as disorders of brain circuits and assumes that the dysfunction in neural circuits can be identified with clinical neuroscience tools for quantifying neural connections in vivo, such as electrophysiology and functional neuroimaging (6). In particular, resting-state functional magnetic resonance imaging (rs-fMRI) is a valuable tool for this purpose because it allows non-invasive and quantitative investigation of whole-brain resting-state functional connectivity (rs-FC) (7, 8, 9). Numerous findings on neural circuit abnormalities in psychiatric disorders using rs-fMRI have been reported (10, 11, 12, 13). Furthermore, several studies have described neurobiologically homogeneous subgroups in psychiatric disorders by objective measures, including rs-FC, rather than relying on conventional clinical diagnosis (14, 15, 16, 17). The research results based on the RDoC concept are expected to create objective biomarkers for stratifying psychiatric disorders in a pathophysiologically uniform manner and eventually to improve the outcomes of psychiatric patients.

Several pioneering studies utilized rs-FC for stratifying specific types of MDD patients into subgroups, and successfully predicted their different responses to a few treatments. Drysdale et al. stratified MDD patients based on biology and behavior and suggested the existence of four distinct subtypes of MDD (18). The authors employed canonical correlation analysis (CCA) to determine a two-dimensional mapping between rs-FC data and depressive symptoms, then applied a hierarchical clustering on the two components derived from CCA and identified four subtypes of MDD patients. Notably, these subtypes were predictive of transcranial magnetic stimulation (TMS) treatment response. Tokuda et al. combined rs-FC with clinical questionnaire scores and various biomarkers in a high-dimensional co-clustering model (19). The model that best distinguished MDD patients from healthy controls identified three subtypes of MDD patients. These MDD subtypes were characterized by differences in rs-FC (between the angular gyrus and the DMN) level and different levels of childhood abuse trauma, and also related to a selective serotonin-reuptake inhibitor (SSRI) treatment response. These research results indicate a possibility of appropriate treatment selection for each subtype based on rs-fMRI stratification of MDD patients who are collectively diagnosed as MDD according to conventional diagnostic criteria.

Although there have been several reports on the usefulness of biomarkers in MDD stratification, problems still need to be solved for practical application. In particular, few studies have demonstrated reproducibility i.e., generalizability of fMRI stratification biomarkers in MDD patients on independent validation cohorts. Dinga et al. attempted to replicate the procedure followed in the Drysdale et al., 2017 study and their findings in a different clinical population of depression (20). Although they found high canonical correlations between rs-FC and clinical symptoms and an optimal three-cluster solution as in the original study, neither canonical correlations nor clusters were statistically significant. As one of the reasons, they pointed out the possibility that overfitting in the CCA procedure due to the large number of variables compared to the number of subjects in the samples led to instability in an independent dataset. Regarding the study of Tokuda et al., 2018, the sample they used was relatively small and collected at one facility. Whether their results are replicated in a dataset from other facilities has not been verified.

The difficulty in generalizability may at least come from measurement and sampling biases inherent to multi-site fMRI data and the overfitting of machine learning algorithms to discovery cohorts with small samples, i.e., a lack of appropriate machine learning algorithms for stratification with relatively small datasets. An unsupervised learning technique with data-driven approaches is a promising method for stratification. However, machine learning algorithms tend to fit the distinctive characteristics of the training dataset, resulting in poor performance on independent data. Neglecting to account for this inherent optimism can be particularly precarious in situations where the number of explanatory variables is far larger than the sample size, such as in the context of the rs-fMRI data (21).

To overcome the above difficulties, our group has been making various efforts and reported a number of research results. Generalizable biomarkers that can be replicated with data collected at independent sites require data collection from multiple sites. We have collected large-scale, multi-site, multi-disorder neuroimaging data from thousands of samples in the Japanese Strategic Research Program for the Promotion of Brain Science (SRPBS) project and published it as a database (22). A unified protocol of SRPBS guarantees anterograde data harmonization across multiple imaging sites. Utilizing a traveling-subject dataset in this database, we demonstrated that site differences are composed of biological sampling bias and engineering measurement bias. We developed a novel retrograde harmonization method that removed only the measurement bias (23). Consequently, imaging data can be both anterogradely and retrogradely harmonized. Furthermore, to suppress overfitting and successfully develop generalizable stratification biomarkers, we proposed the following hierarchical supervised/unsupervised learning strategy (24, Fig. 1).

**Fig. 1.**
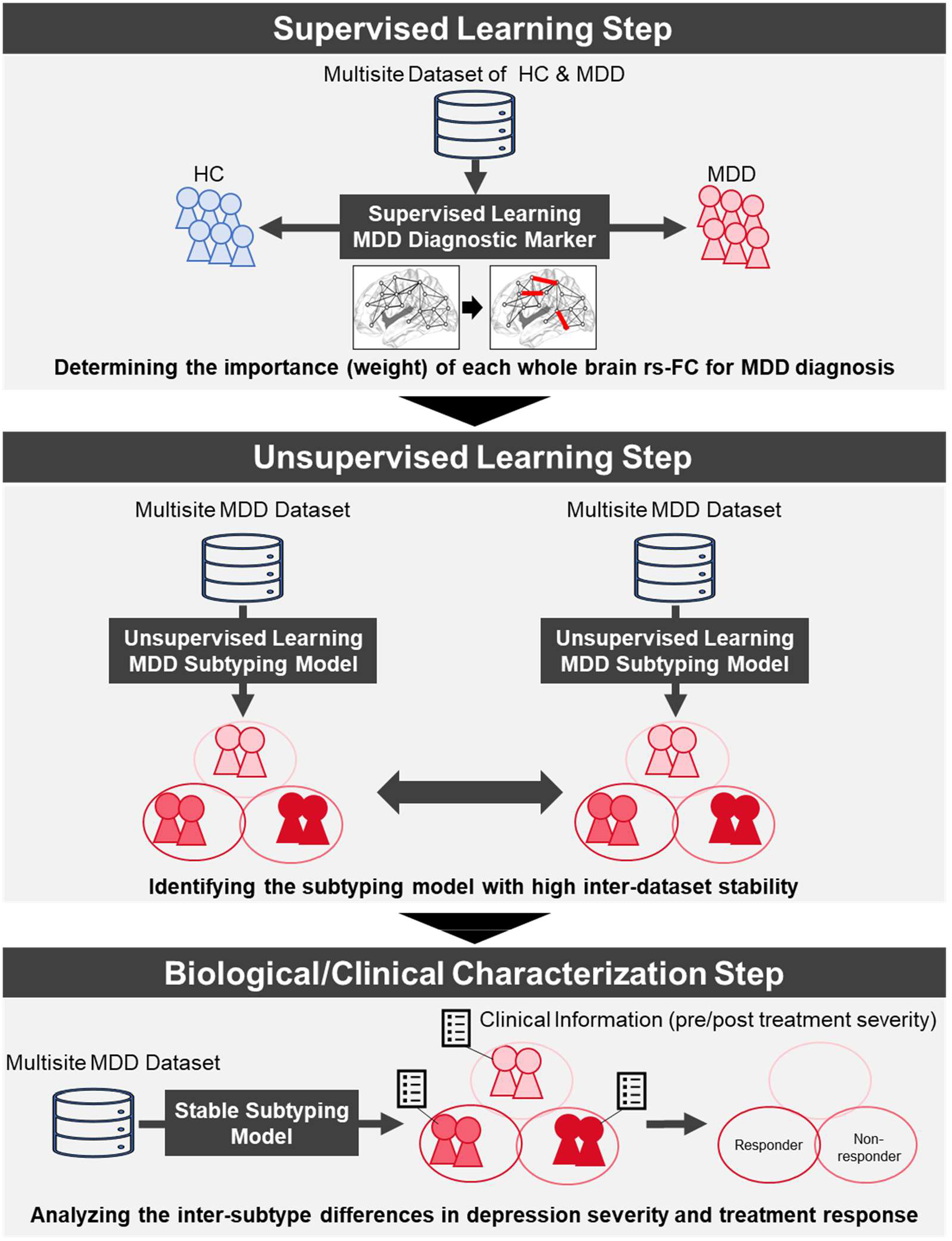
Overview of supervised/unsupervised learning strategy to develop robust MDD stratification biomarkers. In the first step, using supervised learning, MDD diagnostic biomarkers with generalization performance are developed to distinguish MDD patients from healthy controls. Based on the developed MDD diagnostic biomarkers, the importance of each whole-brain rs-FC in distinguishing MDD patients from healthy controls is determined. In the second step, MDD patients are stratified by unsupervised learning using only the highly important rs-FCs determined by the diagnostic biomarkers, and a stratification biomarker (subtyping model) is simultaneously obtained. In this step, unsupervised learning is performed on each of two independent multi-site MDD datasets to evaluate the clustering stability between the datasets. In this study, multiple MDD stratification biomarkers are developed based on various MDD diagnostic biomarkers, and the MDD stratification biomarker with the highest clustering stability is identified. Finally, based on the fMRI data of each MDD patient and the accompanying clinical information, the neurobiological and clinical characteristics, such as treatment responsiveness of each MDD subtype, are clarified.

**Fig. 2.**
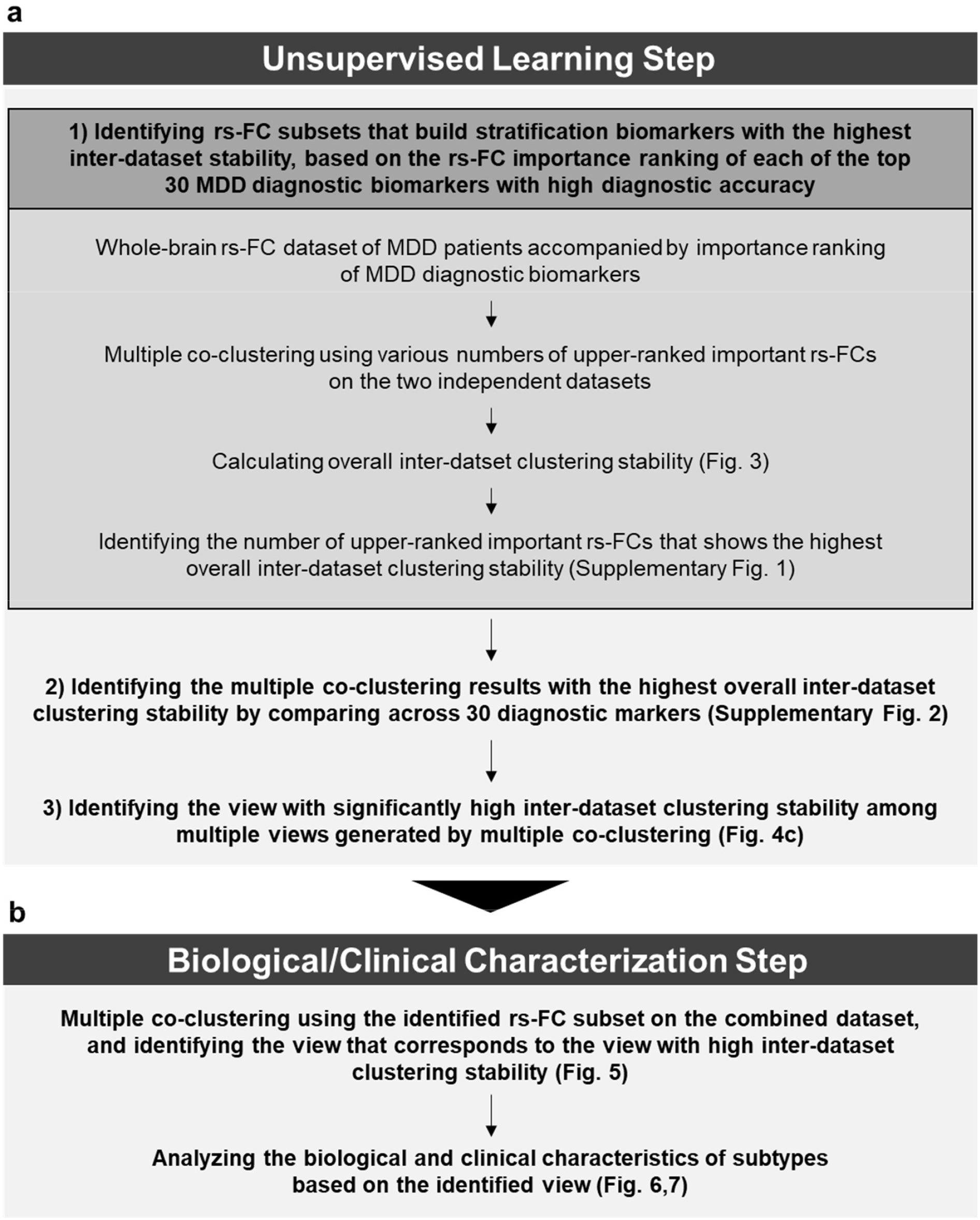
Overall flow of this research

First, using a supervised learning algorithm, we identify a relatively small number of rs-FCs that reliably distinguish healthy controls (HCs) and patients with psychiatric disorders. This stage aims to find potentially relevant biological dimensions to psychiatric disorders. Second, we apply unsupervised learning to these low biological dimensions to find subtypes of psychiatric disorders and identify stratification biomarkers with high inter-dataset stability. Finally, we analyze the clinical characteristics of each subtype using individual patient clinical information accompanying rs-fMRI data. As a first step in this strategy, our group has already successfully developed many diagnostic biomarkers with generalizability through supervised learning for several psychiatric disorders using the above database and harmonization method for site differences (24, 25, 26, 27, 28, 29).

This study aimed to develop generalizable biomarkers for stratifying MDD patients and to elucidate the clinical characteristics, such as treatment responsiveness of the stratified MDD patient subtypes, as the second and final steps in the hierarchical supervised/unsupervised learning strategy. We used the rs-FC subsets with high diagnostic importance determined by the MDD diagnostic biomarker constructed in previous research (29). We employed multiple co-clustering (30) as an unsupervised learning algorithm which is expected to further reduce dimensionality by dividing features and construct generalizable stratification biomarkers. Based on many MDD diagnostic biomarkers, we constructed many MDD stratification biomarkers and identified the stratification biomarker with the highest clustering stability between two independent datasets evaluated using a novel method. Finally, we analyzed the responsiveness of the stratified MDD patient subtypes to a selective serotonin reuptake inhibitor (SSRI) treatment using longitudinal depression severity data accompanying rs-MRI data.

## MATERIALS AND METHODS

### Overview of hierarchical supervised/unsupervised learning strategy

Fig. 1 shows the overview of this study following a hierarchical supervised/unsupervised learning strategy. For the first supervised learning step, i.e., developing MDD diagnostic biomarkers that discriminates between MDD patients and HCs, we utilized the recent achievements of our group (29). A total of 360 development pipelines were implemented by preparing multiple options for each step in the development of diagnostic biomarkers and combining them. As the next step, in this research, unsupervised learning was performed using rs-FCs, which is highly important in the 30 diagnostic biomarkers with the highest discrimination performance (29, Supplementary Table 1). As an unsupervised learning algorithm, we used multiple co-clustering which is helpful for clustering with high-dimensional data and can identify clustering patterns of subjects based on rs-FCs in specific brain areas (30). Using the two independent multiple-site rs-fMRI datasets (Dataset-1 and Dataset-2, Supplementary Table 2), we evaluated the inter-dataset stability of the developed MDD stratification biomarkers and identified the most stable stratification biomarker between the datasets. Finally, to associate the subtyping with the clinical characteristics, we analyzed the treatment response to antidepressants of each subtype.

### Participants

We analyzed the same rs-fMRI datasets (Supplementary Table 2) previously presented in Yamashita et al., 2020 (24) and Takahara et al., 2024 (29): Dataset-1 contained data from 612 participants (463 HCs from four sites and 149 depressive patients from three sites); Dataset-2 contained data from 449 participants (264 HCs and 185 depressive patients from four sites). The 103 depression patient samples out of the 334 in total of Dataset-1 and Dataset-2 have the depression severity scores (Hamilton Rating Scale for Depression: HAMD) before and 6-8 weeks after the onset of SSRI treatment, and that data was used to analyze treatment response. Note that rs-fMRI data for those subjects were obtained before or within two weeks after the onset of treatment with SSRI. Most rs-fMRI data utilized in this study can be downloaded publicly from the DecNef Project Brain Data Repository (https://bicr-resource.atr.jp/srpbsopen/ and https://bicr.atr.jp/dcn/en/download/harmonization/). All participants in the datasets provided written informed consent to participate. All the recruitment procedures and experimental protocols were approved by the respective institutional review boards and conducted under the Declaration of Helsinki.

### Preprocessing and rs-FC computation

Calculating rs-FC from rs-fMRI images was performed according to the previously reported method (24, 29). Briefly, the rs-fMRI data were preprocessed using fMRIPrep version 1.0.8 (35), which consisted of several steps, including the exclusion of the initial 10 seconds of data for T1 equilibration, slice-timing correction, realignment, coregistration, distortion correction using a field map, segmentation of T1-weighted structural images, normalization to the Montreal Neurological Institute space, and surface projection. Due to the lack of field map data, “Fieldmap-less” distortion correction was performed on Dataset-2. Since the coregistration was unsuccessful for the data of six subjects in Dataset-2, we excluded these data from further analysis. We used linear regression with 12 regression parameters: six motion parameters, average signals over the whole brain, and five anatomical CompCor (36) components to remove several sources of spurious variance. To restrict the analysis to low-frequency fluctuations which are characteristic of rs-fMRI BOLD activity (37), a temporal bandpass filter was applied to the time series using a second-order Butterworth filter with a pass band of 0.01–0.08 Hz. A scrubbing procedure was performed using frame displacement (FD) based on head motion. To exclude any volume with excessive head motions, we removed the volumes with FD > 0.5 mm, as proposed in a previous study (38). This threshold removed 6.3% ± 13.5 volumes (mean ± SD) per rs-fMRI session from all the datasets. To keep data quality high enough, the subject data with more than the mean + 3 SD excluded volumes were removed, resulting in 33 subjects being removed for subsequent analyses. As a result of preprocessing, we included 584 participants (446 HCs and 138 MDDs) in Dataset-1 and 438 participants (259 HCs and 179 MDDs) in Dataset-2. The method we used for the parcellation of brain regions, calculation of rs-FC matrix, and harmonization of site differences were detailly described in Takahara et al., 2024 (29). On each site data of both HCs and MDDs, we standardized each rs-FC using its mean and standard deviation after the age factor was regressed out from each rs-FC data.

### Unsupervised learning algorithm (Multiple co-clustering)

On the unsupervised learning step, we used the multiple co-clustering method developed by Tokuda et al. (19, 30). This method performs clustering based on nonparametric Bayesian mixture models in which features are automatically partitioned for each subject clustering solution by maximizing the marginal likelihood. The multiple-subject clustering solutions can be obtained depending on the number of feature partitions. The feature partitioning works as a feature selection for different subject cluster solutions. Here we refer to a co-clustering solution that provides both feature- and subject-clusters as a ‘view’. For each view, the subtyping model which gives the way to cluster new subject data is described by the Gaussian mixture distribution. It should be noted that the subtyping models are trained differently depending on the initial configurations. To obtain the robust subtyping model, we used the random starting points strategy in which we trained the model with multiple different initial configurations, and then selected the most consistent model among models with high likelihood values. Specifically, we used 1000 and 10000 starting points for the inter-dataset clustering stability analysis and the clinical characterization analysis, respectively. The top 20 models based on the marginal likelihood were selected, and the mean clustering similarity between pairs of the 20 models was calculated. Finally, we chose a model with the highest mean clustering similarity.

### Procedure for evaluation of inter-dataset clustering stability

The evaluation procedure of clustering stability between independent datasets is shown in Fig. 3a. In the case of unsupervised learning models, it is not possible to evaluate the predictive accuracy on independent datasets like in supervised learning models, as there is no ground truth. Therefore, we considered evaluating the similarity of subject clustering solutions between datasets as an inter-dataset stability of the subtyping model. However, subject clustering solutions yielded by cluster analysis on two independent datasets could not be directly compared because the subjects are entirely different. Then, we subtyped MDD patients in one dataset using subtyping model yielded by learning from another dataset. We evaluated the similarity between the subject clustering solution and the subtyping result by adjusted rand index (ARI) and vice versa. Furthermore, since several subject clustering solutions and subtyping models (multiple views) are yielded by multiple co-clustering in each dataset, we evaluated similarities of all combination patterns of views between datasets. As a result, two sets of ARI values for the number of all those combinations of multiple views (Fig. 3b). Statistical significances of ARI values were confirmed by permutation tests in which the subtyping results by another dataset’s subtyping model were randomly shuffled and compared with clustering solutions 10000 times. An ARI value was considered statistically significant if it exceeded the top 500th (5%) ARI values in the permutation tests. We calculated the average of the two ARI sets (Fig. 3c), then identified the maximum ARI value for each view in the average ARI set and averaged them to obtain the overall inter-dataset clustering stability (Fig. 3d).

**Fig. 3.**
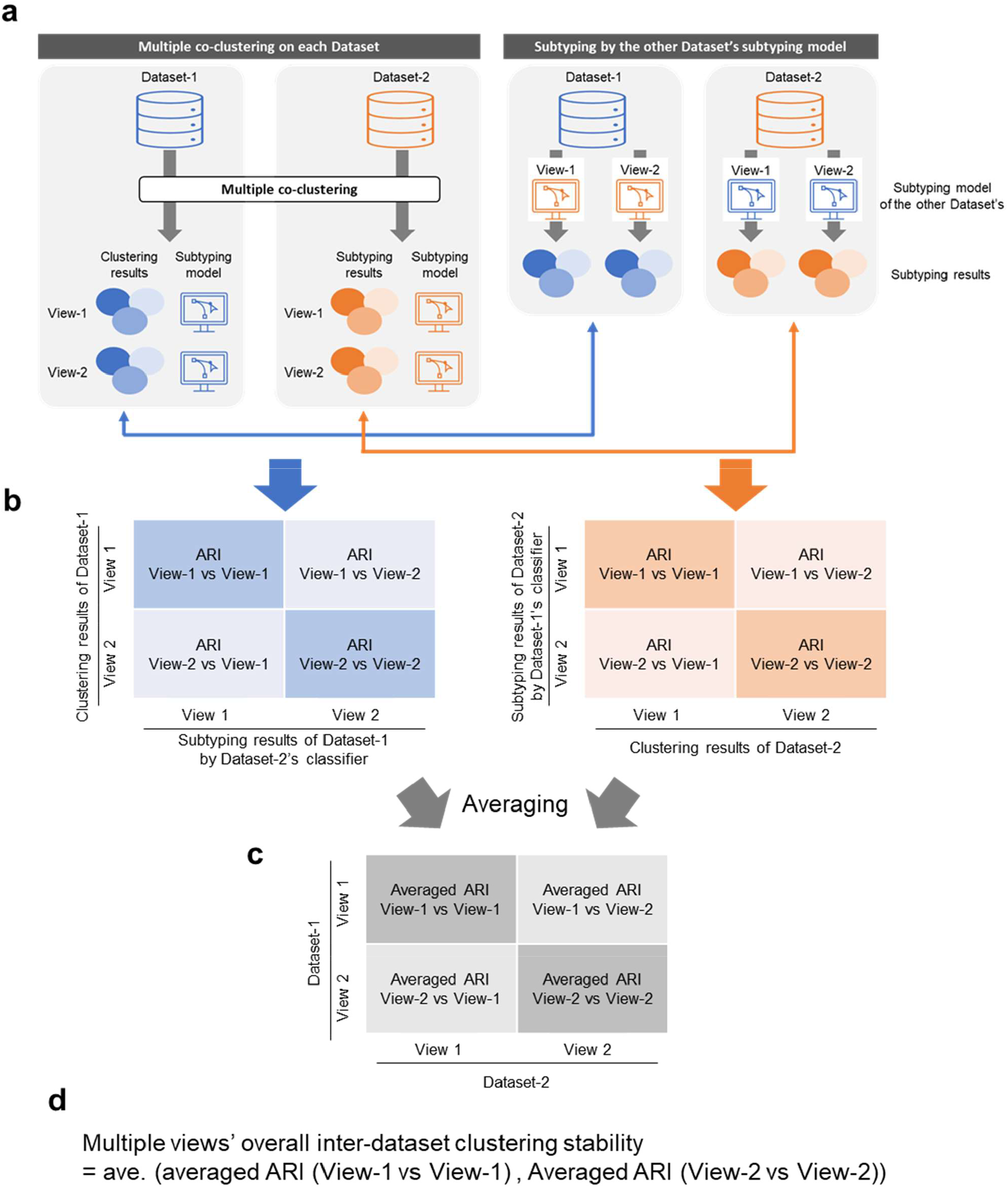
Procedure for assessing clustering stability between independent data sets. a) Multiple views, including clustering results and subtyping models, are generated after performing multiple co-clustering on two independent datasets. Next, the subtyping models of each view are used to subtype MDD patients in the other dataset. b) The degree of agreement (ARI) between the clustering results of MDD patients by multiple co-clustering and the subtyping results using the subtyping model of the other dataset is calculated between all views. c) The ARIs of all inter-view comparisons obtained in each dataset are averaged between the two datasets. d) The overall inter-dataset clustering stability of multiple views is calculated by extracting several corresponding view pairs and averaging the ARIs of those inter-view comparisons.

### Experimental identification of rs-FCs for construction of stable stratification biomarker

We experimentally investigated how many top-ranked rs-FCs in MDD diagnostic biomarkers could construct MDD stratification biomarkers with the highest overall inter-dataset stability. We prepared ten subsample datasets to evaluate dataset fluctuations by excluding 1/10 of data in each dataset and evaluated the overall inter-dataset stability for 100 subsample pairs. In the first search, clustering was performed using the top 50, 100, 200, 300, and 400 rs-FCs, and the overall inter-dataset stability of each was evaluated. In the second search, clustering was performed at intervals of 10 rs-FCs around the number that showed the highest stability in the first search. Then, the number of rs-FCs that showed the highest stability in the second search was identified (Supplementary Fig. 1). To obtain the final results for each MDD diagnostic biomarker, stratification biomarkers were constructed on full-sample of each dataset using the identified number of rs-FCs and the inter-dataset stability was calculated.

### Analysis of the association between subtype and antidepressant treatment response

For the clinical characterization of subtypes, we focused on responses to antidepressant treatment. Due to the limited subject sample with the longitudinal depression severity data necessary for the analysis of treatment response, i.e., depression severity information before and after antidepressant treatment, clustering was performed for all data, including Dataset-1 and Dataset-2 so that the sample size of each subtype was increased for statistical power. From the clustering results, the subject samples with longitudinal depression severity data were extracted, and statistically analyzed the relationship between treatment response and subtyping. The improvement rate of depression severity was calculated by (HAMD before treatment – HAMD after treatment) / HAMD before treatment. Differences in the HAMD score before/after treatment and the improvement rate among subtypes were statistically analyzed by non-parametric ANOVA (Kruskal-Wallis test) and post hoc pairwise non-parametric multiple comparison (Steel-Dwass). The rate of responders and remitters were also evaluated as indices of treatment response. Responders are patients with >50% improvement in HAMD, and remitters are patients whose HAMD has decreased to <8 after treatment. The chi-square test was performed to confirm statistically significant differences in the rate of responders and remitters among subtypes.

## RESULTS

### Identification of the stratification biomarker with the highest inter-dataset stability

For learning the stratification biomarkers using fMRI data of depressed patients, we used two multicenter datasets, which had significantly different facilities and measurement protocols (Dataset-1: a dataset of 138 depressed patients acquired with unified measurement protocols across three facilities; Dataset-2: a dataset of 181 depressed patients acquired with non-unified measurement protocols across four facilities) (24, 29). Stratification biomarkers that demonstrate high stability between datasets with significantly different facilities and measurement protocols are considered to be biomarkers with high generalization performance to diverse data.

Starting from several MDD diagnostic biomarkers constructed in previous studies, the process of identifying one MDD stratification biomarker (View) based on the inter-dataset clustering stability is shown below (Fig.2a).

1. For each of the top 30 diagnostic biomarkers with the highest diagnostic accuracy (Supplementary Table 1), we identified the number of rs-FCs that would provide the highest overall inter-dataset clustering stability (Supplementary Fig. 1).
2. We identified the rs-FC subset (determined by the number of top-ranked rs-FC and the importance ranking of the diagnostic biomarkers) that resulted in the highest overall inter-dataset clustering stability among the 30 diagnostic biomarkers (Supplementary Fig. 2).
3. From the multiple views as an identified result of multiple co-clustering, the view with the statistical significance and the highest stability was identified (Fig. 4c).

**Fig. 4.**
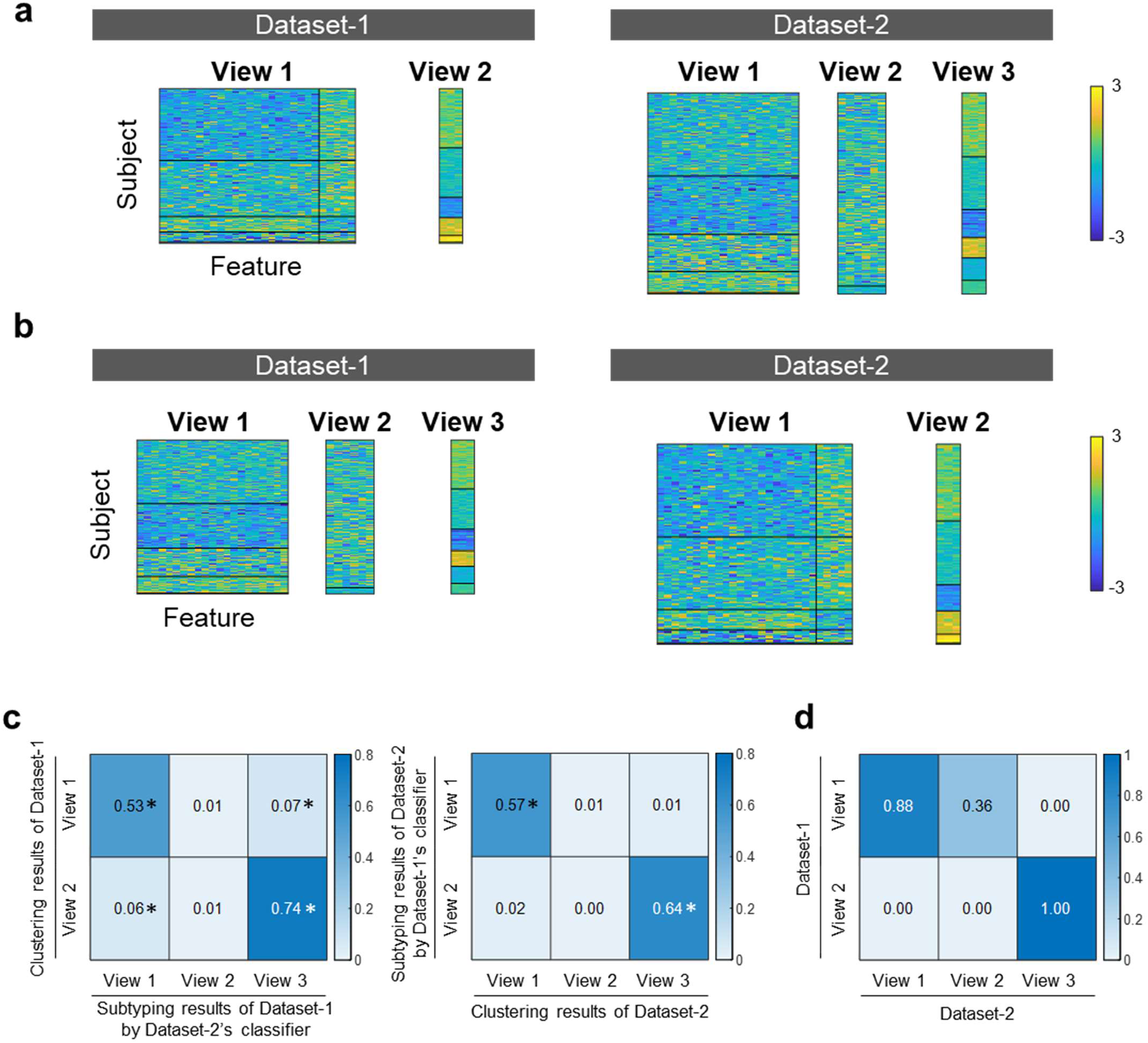
Multiple co-clustering results showing the highest inter-dataset clustering stability. a) Results of multiple co-clustering performed on two independent datasets (visualization of all views). The horizontal axis indicates features, the vertical axis indicates subjects, and the color of each matrix indicates the standardized rs-FC value. b) Results of subtyping each dataset using the subtyping model of the other dataset. c) Evaluation results of clustering stability between datasets. We evaluated the agreement (ARI) between the subject clustering results and the subtyping results using the subtyping model of the other dataset. As there were two views in Dataset-1 and three views in Dataset-2, the ARI evaluated between all views is shown. * indicates that the ARI was confirmed to be a significant value in the permutation test. d) Inter-dataset agreement of rs-FC view assignment. The agreement of rs-FC assigned to each view in each dataset was evaluated using the Dice coefficient.

As a result of steps 1) and 2), the highest overall inter-dataset clustering stability was observed in multiple co-clustering using the 30 top-ranked rs-FCs of the diagnostic biomarker #177 (Supplementary Fig. 1, 2) constructed using BSA for parcellation, Pearson correlation for rs-FC calculation, and Random Forest for supervised machine learning (Supplementary table 1). At this point, we were able to narrow down the whole-brain rs-FCs from thousands to tens of thousands depending on the parcellation to 30 rs-FCs that provided the result of multiple co-clustering with high inter-dataset clustering stability.

As a result of multiple co-clustering using the 30 top-ranked rs-FCs of the diagnostic biomarkers #177, two views were generated in Dataset-1 and three views were generated in Dataset-2 (Fig. 4a). The subtyping results of each dataset using the subtyping model of the other dataset are shown in Fig. 4b. The degree of agreement between the clustering results of the subjects and the subtyping results using the other dataset’s subtyping model of each dataset was examined for all pairs of views and presented as ARI (Fig. 4c).

As a result of the above process 3), among all six pairs of views between the two datasets, two pairs of views showed significantly high ARI, with a particularly high ARI observed between View 2 of Dataset-1 and View 3 of Dataset-2 (Supplementary Table 3 for the cross tables of the view pair). When comparing the View assignments of the rs-FC, the Dice coefficient between View 2 of Dataset-1 and View 3 of Dataset-2, which showed the highest ARI, was 1 (Fig. 4d), which means the three rs-FCs assigned in those views are exactly same.

Even though the two independent datasets were trained separately, the same three rs-FCs were automatically selected by multiple co-clustering, and stable stratification was achieved. This suggests that dividing rs-FCs into multiple views by multiple co-clustering could contribute to the high stability of the stratification biomarkers by further reducing the feature dimensionality.

### Subtypes’ Biological/Clinical Characteristics

We performed multiple co-clustering using the 30 top-ranked rs-FCs of the diagnostic biomarkers #177 on a combined dataset of two independent datasets (Fig. 2b), and two views were generated (Fig. 5b). When we evaluated the degree of agreement with the subject clustering of each dataset (Fig. 5a), View 2 of the combined dataset showed high values with View 2 of Dataset-1 and View 3 of Dataset-2 (Fig. 5c). Furthermore, the rs-FCs assigned to View 2 were the same three connections between the thalamus and postcentral gyrus as those assigned to View 2 of Dataset-1 and View 3 of Dataset-2. View 2 of Dataset-1 and View 3 of Dataset-2 have been shown to have high inter-dataset clustering stability, and View 2 of the combined dataset also exhibited high consistency with those views and utilized the same rs-FC, which indicates that View 2 of the combined dataset can be considered to be a stratification biomarker with high inter-dataset stability that retains the characteristics of View 2 of Dataset-1 and View 3 of Dataset-2.

**Fig. 5.**
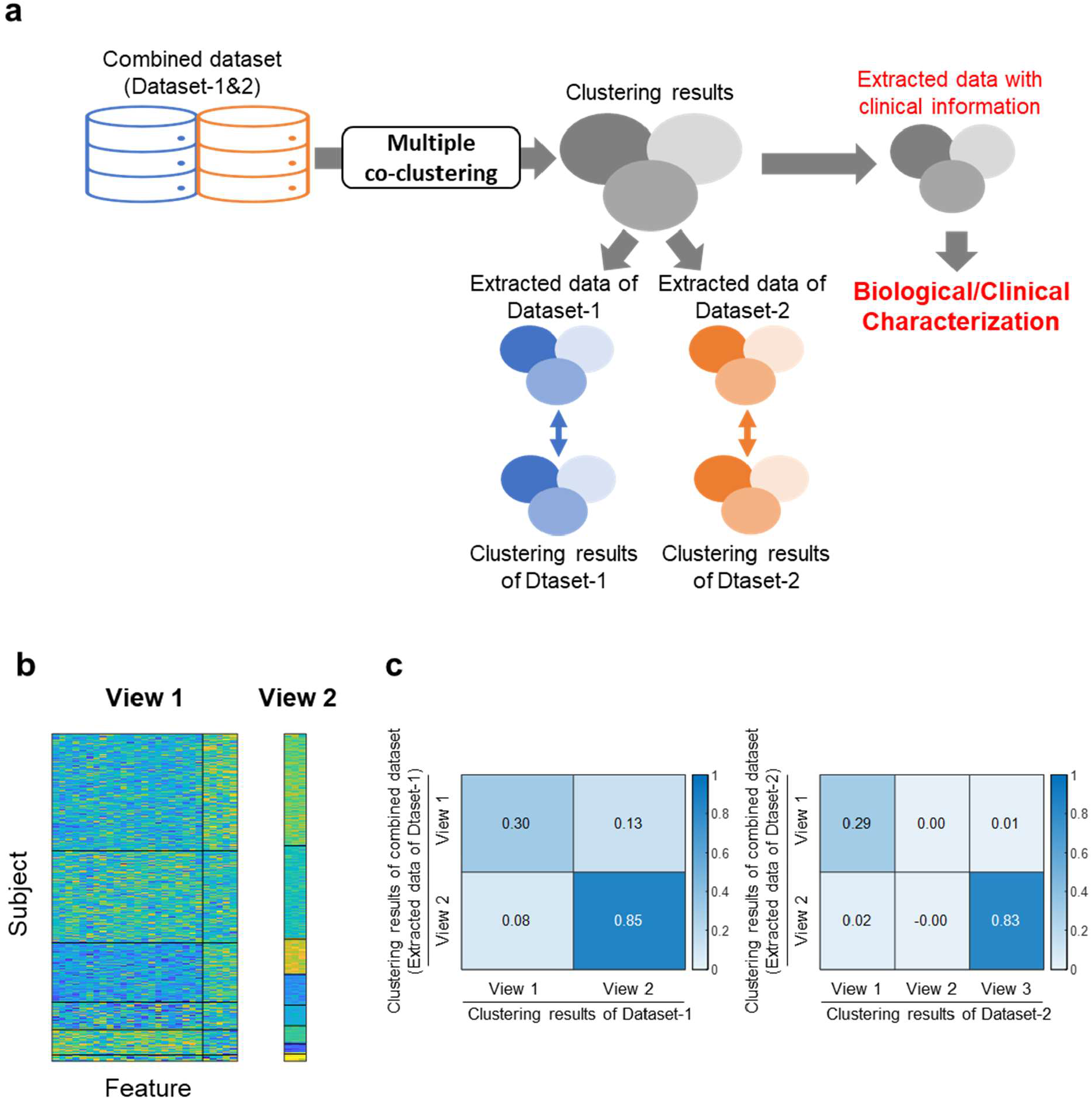
Results of multiple co-clustering in combined datasets for biological/clinical characteristics analysis of subtypes. a) Schema of the neuroscientific/clinical feature analysis of MDD subtypes. To increase the sample size of each subtype, multiple co-clustering was performed on the combined dataset of Dataset-1 and Dataset-2, and views corresponding to View 2 in Dataset-1 and View 3 in Dataset-2, which showed high clustering stability between datasets, were identified. For the clustering results of the identified views, samples accompanied by clinical data were extracted and clinical characteristics were analyzed by comparing subtypes. b) Results of multiple co-clustering of the combined dataset. The horizontal axis indicates features, the vertical axis indicates subjects, and the color of each matrix indicates the standardized rs-FC value. c) The degree of agreement between the subject clustering results of the two views in the combined dataset and the subject clustering results of each view in each dataset.

Fig. 6a shows the three rs-FCs between the thalamus and postcentral gyrus used in the MDD stratification biomarker (View2) with high inter-dataset clustering stability. Since multiple co-clustering is an extension of the Gaussian mixture model, the subject distribution was visualized by a two-dimensional plot of the mean and standard deviation of the three rs-FCs. (Fig. 6b). This figure shows that subjects are primarily clustered based on the average values (horizontal axis) of the three rs-FCs, representing the connectivity strength. To examine the characteristics of these rs-FC strengths for each subtype, the average values of the non-standardized rs-FC values for each subtype were shown in Fig. 6c. Subtype 2 showed a distribution of average standardized rs-FC values around zero (Fig. 6b) and non-standardized rs-FC values close to zero as well (Fig. 6c), indicating that a neurobiological characteristic of this subtype is that the strengths of these rs-FCs are the weakest among all subtypes and have average strength across all subjects. Additionally, it can be observed that the strengths of the rs-FCs were positively stronger in the order of Subtype 8, 3, 1, and 6, and negatively stronger in the order of Subtype 7, 4, and 5 (Fig. 6c).

**Fig. 6.**
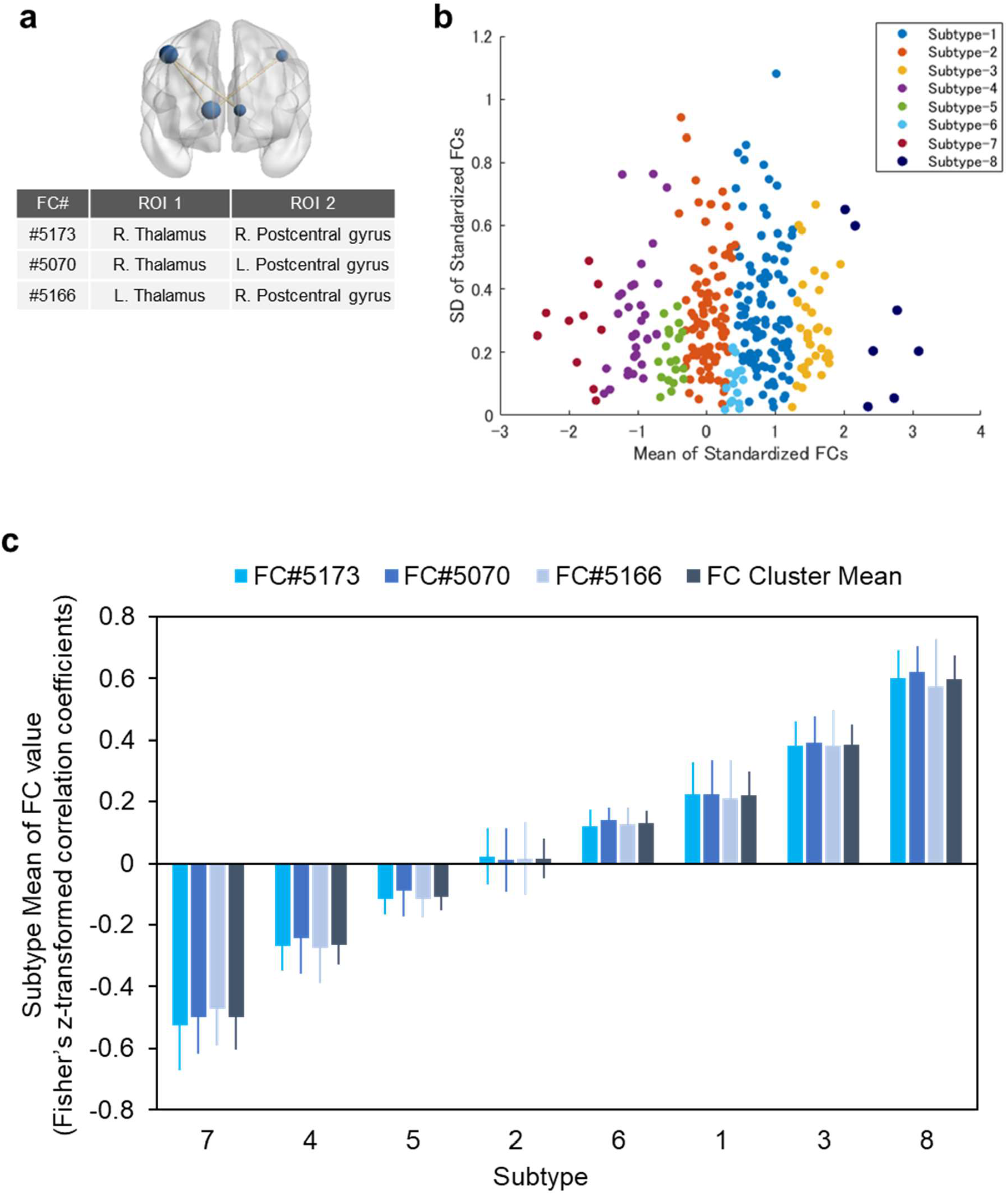
Neurobiological characteristics of MDD subtypes. a) The three rs-FCs used in stratification biomarkers were identified based on clustering stability between datasets. b) Visualization of the subject distribution of the three rs-FCs. The horizontal axis shows the average of the three rs-FCs, and the vertical axis shows the standard deviation. The rs-FC values were standardized using subject data from each facility. c) The average of the three rs-FCs and their averages for each subtype. To confirm the strength of the rs-FCs, Fisher’s z-transformed correlation coefficients before standardization are shown.

An analysis of clinical characteristics was conducted on Subtype 1, 2, and 4 which included at least ten subjects with longitudinal depression severity data (Fig. 7a). These subtypes were labeled based on the neurobiological characteristics of the three rs-FCs’ strengths as follows: Subtype 1, Positive-correlation subtype; Subtype 2, Weak-correlation subtype; Subtype 4, Negative-correlation subtype. To analyze the treatment responsiveness of each subtype, the depression severity (HAMD score) at the initiation of SSRI treatment and after 6-8 weeks, severity improvement rate, response rate, and remission rate were shown in Fig. 7b-7e. No significant differences in baseline severity were observed among the three subtypes (P = 0.727). In contrast, significant differences were found in the depression severity after 6-8 weeks of treatment (P = 0.014), with the Weak-correlation subtype having significantly lower severity compared to the Negative-correlation subtype (P = 0.026). A significant difference was also observed in the severity improvement rate after 6-8 weeks of treatment among the subtypes (P = 0.003), with the Weak-correlation subtype showing significantly higher improvement rates compared to both the Negative-correlation subtype (P = 0.011) and the Positive-correlation subtype (P = 0.020). Significant differences were found in the response rate and remission rate among the subtypes as well (P = 0.032 for responder rate; P = 0.014 for remission rate), with the Weak-correlation subtype showing significantly higher rates compared to the Negative-correlation subtype (P = 0.030 for responder rate; P = 0.018 for remission rate). Moreover, when analyzing all subtypes, including those with less than ten subjects, a significant association between subtyping and treatment responsiveness was observed (Supplementary Fig. 4). On the other hand, no significant association was found between confounding factors such as age, gender, and imaging facility and subtyping (Supplementary Fig. 5). Since significant differences in treatment response were observed between subtypes, we confirmed the relationship between the connectivity strength of the thalamus-postcentral gyrus, a neurobiological characteristic of the subtype, and treatment response (Fig. 7f). MDD patients with standardized average rs-FC strengths near 0, indicating strengths close to the mean across all subjects, showed higher improvement rates (Fig. 7f). Conversely, MDD patients with positively or negatively strong rs-FC strengths tended to have lower improvement rates.

**Fig. 7.**
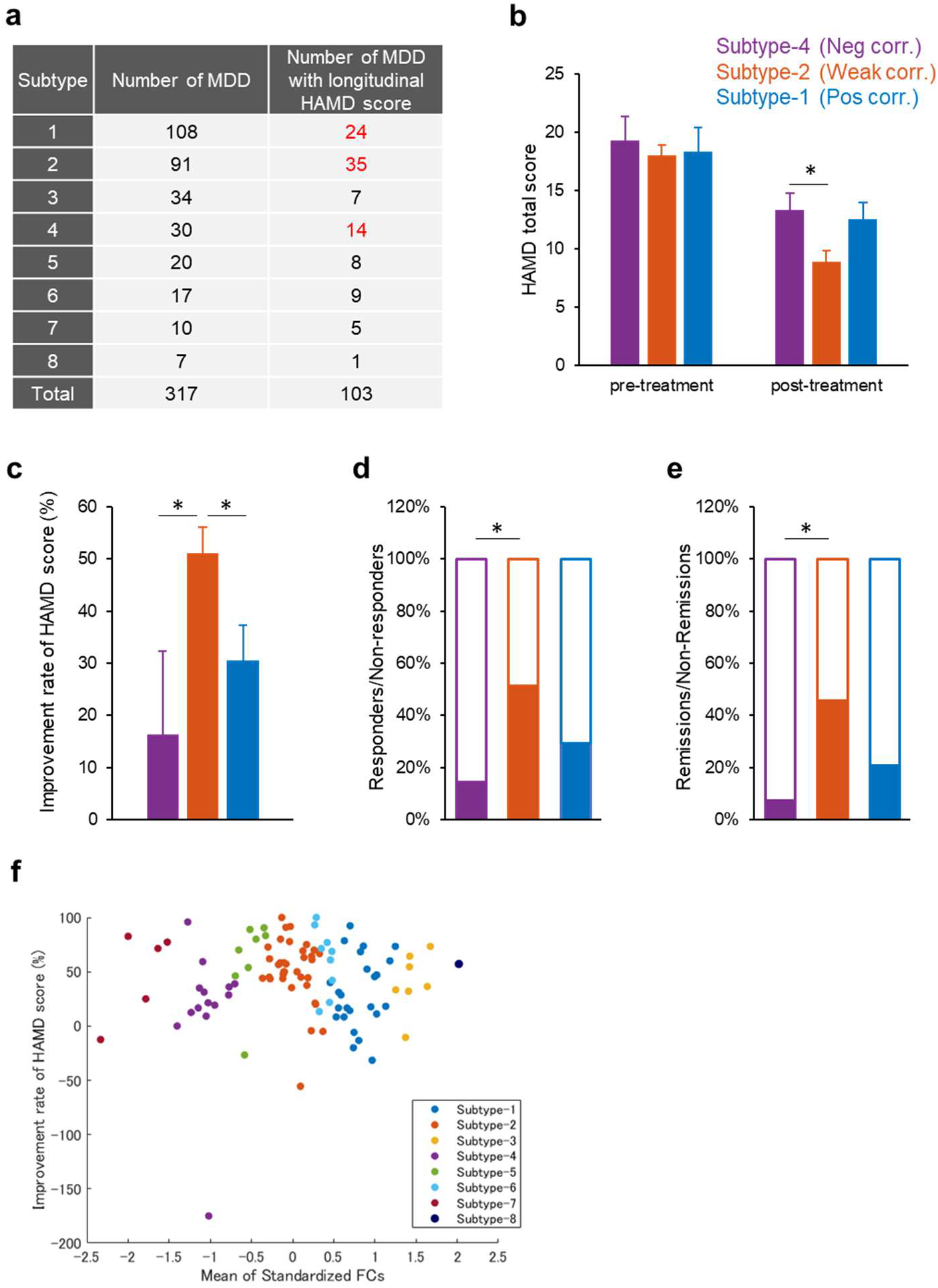
Clinical characteristics of MDD subtypes. The clinical characteristics of the subtypes were analyzed with a focus on the therapeutic response to antidepressants. a) The number of MDD patients in each subtype, and the number of MDD patients with HAMD score before and 6-8 weeks after the onset of SSRI treatment which are necessary for the analysis of therapeutic response. Subtypes 1, 2, and 4 which had at least 10 MDD patients with longitudinal HAMD scores (shown in red) were analyzed. b) The mean HAMD score of each subtype before and 6-8 weeks after the onset of treatment. *P<0.05 by Steel-Dwass test following Kruskal-Wallis test. c) HAMD improvement rate of each subtype. HAMD improvement rate = (HAMD score before treatment - HAMD score 6-8 weeks after the onset of treatment) / HAMD score before treatment *100. *P<0.05 by Steel-Dwass test following Kruskal-Wallis test. d) Response rate indicates percentage of subjects showing >50% reduction in HAMD score. *P<0.05: Tested by chi-square test (FDR corrected). e) Remission rate indicates percentage of subjects whose post-treatment HAMD score fell below 8 points. *P<0.05: Tested by chi-square test (FDR corrected). f) Relationship between the three rs-FCs used as stratification biomarkers and treatment response. The horizontal axis shows the standardized average value of the three rs-FCs, and the vertical axis shows the HAMD improvement rate.

### Multiple co-clustering with clinical information

Finally, we examined whether stratification related to treatment response with SSRI could be achieved using multiple co-clustering based on clinical evaluation indices (Supplementary Fig. 6). Similar to the above when analyzing subtypes that included at least ten subjects, significant differences in depression severity were observed among subtypes at the beginning of treatment (P < 0.001), as well as at 6-8 weeks after the start of treatment (P < 0.001). However, no significant differences in severity improvement rates were observed among subtypes (P = 0.085). These results suggest that stratifying MDD patients based on biological features such as rs-FC, rather than classifying them based on clinical observations like severity, may be more effective in predicting treatment outcomes.

## DISCUSSION

In this study, we succeeded in developing a biomarker that allows stable stratification between entirely independent datasets. Furthermore, since the responsivity to antidepressants differs among several MDD subtypes, this stratification biomarker may help predict treatment response to antidepressants.

An essential issue for biomarkers created using machine learning technology is generalization performance, that is, whether the performance can be reproduced even with data different from the learning data. Unlike diagnostic biomarkers created using supervised learning, there is no established method for evaluating generalization performance for stratification biomarkers created using unsupervised learning because there is no true answer. Drysdale et al. showed that diagnostic biomarkers created by supervised learning for each MDD biotype stratified by unsupervised learning generalized to independent data, but the generalization performance of the stratification biomarker itself was not evaluated (18). Dinga et al. evaluated the stability of the stratification biomarker of Drysdale et al. (18) by removing one subject from the training data, but this evaluation method is highly conservative (20). We have devised a unique method to assess the stratification stability, i.e. the generalizability performance of stratification biomarkers, across entirely independent datasets. In this evaluation method which is much more stringent than Dinga et al.’s conservative method, our stratification biomarker showed high stability across entirely independent datasets, indicating high generalizability performance.

We believe that there are two main reasons for our success in developing a stratification biomarker with high generalizability. The first is the dataset used for learning. Our group has placed importance on the quantity and quality of data in developing biomarkers with high generalizability. We have been collecting rs-fMRI data from patients with psychiatric disorders at multi-site in large-scale research projects (22). Although many fMRI biomarker studies used data from dozens of subjects acquired at one institution, the two datasets used in this study both consisted of rs-fMRI data from over 100 MDD patients acquired at 3-4 multiple institutions. The second is the strategy for developing stratification biomarkers. We believe that selecting the features used for clustering would be an important key to create stable stratification biomarkers. Dinga et al. pointed out that in Drysdale et al.’s study (18), information leakage during the feature generation process using CCA was the cause of the biomarkers being overfitted to the training data and having low stability (20). In our supervised/unsupervised learning strategy, we first created a depression diagnostic biomarker by supervised learning using the whole-brain rs-FCs. Then, we used rs-FCs which are highly important in distinguishing between depressed patients and healthy subjects, as features in unsupervised learning to create a stratification biomarker. Furthermore, we used multiple co-clustering for the unsupervised learning algorithm, which avoids overfitting by appropriately grouping many features and allows for more reliable clustering. The collection and utilization of high-quality and quantity datasets and the supervised/unsupervised learning strategy have led to the success of developing highly stable stratification biomarkers.

Significant differences in depression improvement rates with antidepressant treatment were observed among the three MDD patient subtypes identified by the stratification biomarker. The rs-FCs used in this subtyping model connect the thalamus and the postcentral gyrus in the sensorimotor area. Several reports showed an association of the thalamus with depression pathophysiology and treatment response. Yamamura et al. reported that depressed patients with treatment resistance had altered neural activity in the thalamus, and its activity correlated with antidepressant treatment response (31). The sensorimotor cortical regions are structurally and functionally interconnected with the thalamus (32). The connectivity between the thalamus and sensorimotor network (SMN) has been reported to be associated with psychomotor abnormality in psychiatric disorders (33). Martino et al., reported that decreased thalamo-SMN FC was observed in depressed BD patients with psychomotor retardation, whereas patients with mania exhibited almost the opposite pattern of increased thalamo-SMN FC (34). In our results, subjects with near-average thalamic-postcentral gyrus rs-FC were highly responsive to treatment, whereas subjects with higher or lower rs-FC showed a trend of low treatment response. The rs-FC and treatment response appear to have an inverse U-shaped relationship (Fig. 7f). Northoff et al. proposed that the relationship between neural measurements and mental function can be characterized by a continuous nonlinear inverted-U shaped curve (33). In this relationship, the average state of a neural measure is associated with optimal mental function, whereas extreme states, either high or low, are associated with unfavorable mental function. They summarized this model of the neuro-mental relationship as “average is good, extremes are bad”. The inverted U-shaped relationship observed between the thalamic-postcentral gyrus rs-FC and treatment response in this study consists with the model of Nortoff et al.

In this study, more than 300 MDD patients were stratified using fMRI data, but 103 had longitudinal depression severity data to analyze the treatment response of subtypes. Although it was confirmed that the clustering structure of the subset of samples was not significantly different from the clustering structure of all patient samples (Supplementary Fig. 3), it was not possible to clarify the clinical characteristics of all subtypes because the subtypes with a small number of samples (less than ten patients) were not included in the analysis, to ensure the reliability of the statistical analysis. However, a preliminary analysis of all subtypes revealed a statistically significant relationship between subtype and treatment response (Supplementary Fig. 4). Based on these results, it is expected that the clinical characteristics of all subtypes would be clarified by accumulating data and increasing the number of samples of each subtype. To clarify the scientific validity of this stratification biomarker, it is also essential to confirm the reproducibility of newly collected data. Furthermore, The MDD stratification biomarker may be related to treatment responsiveness not only to SSRI but also to other depression treatments, because this stratification biomarker was not constructed in a way linked to a specific treatment, but was created through unsupervised learning using only rs-FCs information. To address these research questions, we are currently conducting a clinical study to obtain rs-fMRI data and longitudinal clinical information on depressed patients who received SSRIs or various other treatments. Suppose we can further advance our research and put this stratification biomarker into practical use. In that case, it is expected that we will be able to solve the central problem of low remission rates in MDD.

## Acknowledgments

This study was mainly supported by the collaboration fund of XNef, Inc. and SHIONOGI & CO., LTD. The data resource was supported by JSPS KAKENHI Grant Number JP20H03605, JP21H05174 and JP21H05171, AMED under Grant Number JP22dm0307008, JP22dm0307009, JP19dm0207069, JP18dm0307001 and JP18dm0307004, Moonshot R&D Grant Number JPMJMS2021, and by UTokyo Institute for Diversity and Adaptation of Human Mind (UTIDAHM), and the International Research Center for Neurointelligence (WPI-IRCN) at The University of Tokyo Institutes for Advanced Study (UTIAS).

