## Supplemental Materials for "Prediction of antidepressant treatment response with thalamo-somatomotor functional connectivity revealed by generalizable stratification of depressed patients"

**Supplementary Table 1. Pipelines for constructing MDD diagnostic biomarkers with the top 30 diagnostic performance developed in previous study (29).**

| Ranking | Pipeline ID | Parcellation | FC | Harmonization | Machine learning |
|---|---|---|---|---|---|
| 1 | #33 | Glasser + 19 ROIs (379) | Tangent | No Correction | SVM |
| 2 | #31 | Glasser + 19 ROIs (379) | Tangent | No Correction | RIDGE |
| 3 | #9 | Glasser + 19 ROIs (379) | Pearson | ComBat | Random Forest |
| 4 | #123 | Shen (268) | Pearson | No Correction | Random Forest |
| 5 | #249 | Dictionary Learning (80) | Tangent | No Correction | SVM |
| 6 | #299 | FIND lab (78) | Tangent | No Correction | SVM |
| 7 | #247 | Dictionary Learning (80) | Tangent | No Correction | RIDGE |
| 8 | #15 | Glasser + 19 ROIs (379) | Pearson | No Correction | Random Forest |
| 9 | #225 | Dictionary Learning (80) | Pearson | ComBat | Random Forest |
| 10 | #16 | Glasser + 19 ROIs (379) | Pearson | No Correction | SVM |
| 11 | #171 | BSA atlas (145) | Pearson | ComBat | Random Forest |
| 12 | #14 | Glasser + 19 ROIs (379) | Pearson | No Correction | RIDGE |
| 13 | #231 | Dictionary Learning (80) | Pearson | No Correction | Random Forest |
| 14 | #117 | Shen (268) | Pearson | ComBat | Random Forest |
| 15 | #248 | Dictionary Learning (80) | Tangent | No Correction | Random Forest |
| 16 | #13 | Glasser + 19 ROIs (379) | Pearson | No Correction | LASSO |
| 17 | #124 | Shen (268) | Pearson | No Correction | SVM |
| 18 | #121 | Shen (268) | Pearson | No Correction | LASSO |
| 19 | #63 | Glasser (360) | Pearson | ComBat | Random Forest |
| 20 | #243 | Dictionary Learning (80) | Tangent | ComBat | SVM |
| 21 | #298 | FIND lab (78) | Tangent | No Correction | Random Forest |
| 22 | #278 | FIND lab (78) | Pearson | ComBat | Random Forest |
| 23 | #172 | BSA atlas (145) | Pearson | ComBat | SVM |
| 24 | #195 | BSA atlas (145) | Tangent | No Correction | SVM |
| 25 | #283 | FIND lab (78) | Pearson | No Correction | Random Forest |
| 26 | #226 | Dictionary Learning (80) | Pearson | ComBat | SVM |
| 27 | #177 | BSA atlas (145) | Pearson | No Correction | Random Forest |
| 28 | #8 | Glasser + 19 ROIs (379) | Pearson | ComBat | RIDGE |
| 29 | #26 | Glasser + 19 ROIs (379) | Tangent | ComBat | Random Forest |
| 30 | #279 | FIND lab (78) | Pearson | ComBat | SVM |

**Supplementary Table 2. Demographic information of participants.**

| | Site | HC | | | | MDD | | | | Number of MDD with longitudinal HAMD score |
|---|---|---|---|---|---|---|---|---|---|---|
| | | Number | Male/Female | Age (y) | BDI | Number | Male/Female | Age (y) | BDI | |
| Dataset-1 | COI | 124 | 46/78 | 51.9 ± 13.4 | 8.2 ± 6.3 | 70 | 31/39 | 45.0 ± 12.5 | 26.2 ± 9.9 | 28 |
| | KUT | 169 | 100/69 | 35.9 ± 13.6 | 6.0 ± 5.4 | 17 | 11/6 | 43.9 ± 13.3 | 27.7 ± 10.1 | 0 |
| | UTO | 170 | 78/92 | 35.6 ± 17.5 | 6.7 ± 6.5 | 62 | 36/26 | 38.7 ± 11.6 | 20.4 ± 11.4 | 0 |
| Dataset-2 | HKH | 29 | 12/17 | 45.4 ± 9.5 | 5.1 ± 4.6 | 33 | 20/13 | 44.8 ± 11.5 | 28.5 ± 8.7 | 17 |
| | HRC | 49 | 13/36 | 41.7 ± 11.7 | 9.1 ± 8.5 | 16 | 6/10 | 40.5 ± 11.5 | 35.3 ± 9.5 | 6 |
| | HUH | 66 | 29/37 | 34.6 ± 13.0 | 6.9 ± 5.9 | 57 | 32/25 | 43.3 ± 12.2 | 30.9 ± 9.0 | 52 |
| | UYA | 120 | 50/70 | 45.9 ± 19.5 | 7.1 ± 5.6 | 79 | 36/43 | 50.3 ± 13.6 | 29.7 ± 10.7 | 0 |

HC, healthy control; MDD, major depressive disorder; BDI, Beck Depression Inventory-II; COI, Center of Innovation in Hiroshima University; KUT, Kyoto University; UTO, University of Tokyo; HKH, Hiroshima Kajikawa Hospital; HRC, Hiroshima Rehabilitation Center; HUH, Hiroshima University Hospital; UYA, Yamaguchi University.

**Supplementary Table 3. Cross table between View 2 of Dataset-1 and View 3 of Dataset-2, where the highest clustering stability was observed.**

| | | Dataset-1 subtyping by View-3 of Dataset-2 subtype classifier | | | | | |
|---|---|---|---|---|---|---|---|
| | Subtype # | 1 | 2 | 3 | 4 | 5 | 6 |
| View-2 of Dataset-1 clustering solution | 1 | 45 | 6 | 0 | 0 | 0 | 2 |
| | 2 | 0 | 40 | 0 | 0 | 4 | 0 |
| | 3 | 0 | 0 | 16 | 0 | 2 | 0 |
| | 4 | 0 | 0 | 0 | 16 | 0 | 0 |
| | 5 | 0 | 0 | 0 | 6 | 0 | 0 |
| | 6 | 1 | 0 | 0 | 0 | 0 | 0 |

| | | Dataset-2 subtyping by View-2 of Dataset-1 subtype classifier | | | | | |
|---|---|---|---|---|---|---|---|
| | Subtype # | 1 | 2 | 3 | 4 | 5 | 6 |
| View-3 of Dataset-2 clustering solution | 1 | 55 | 0 | 0 | 2 | 0 | 0 |
| | 2 | 3 | 44 | 0 | 0 | 0 | 0 |
| | 3 | 0 | 2 | 23 | 0 | 0 | 0 |
| | 4 | 0 | 0 | 0 | 17 | 1 | 0 |
| | 5 | 0 | 18 | 2 | 0 | 0 | 0 |
| | 6 | 11 | 1 | 0 | 0 | 0 | 0 |

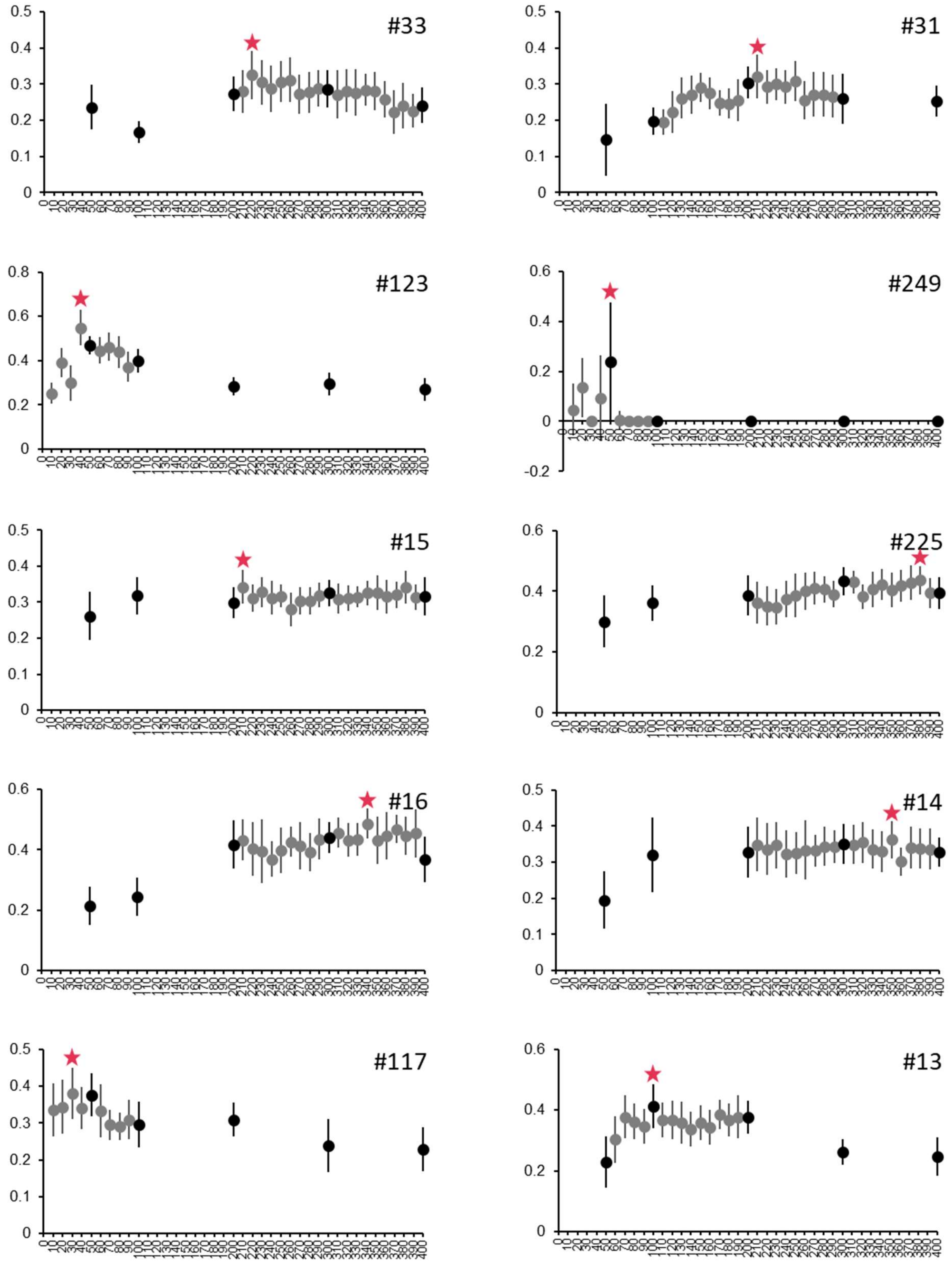

**Supplementary Fig. 1 Overall inter-dataset clustering stability of multiple co-clustering using top-ranked rs-FCs of each MDD diagnostic biomarker.** The horizontal axis indicates the number of top-ranked rs-FCs of MDD diagnostic biomarkers used in multiple co-clustering, and the vertical axis indicates the ARI value. The numbers in the upper right corner of each plot indicate the MDD diagnostic biomarker construction pipeline IDs in Supplementary Table 1. The star marks indicate the highest ARI values.

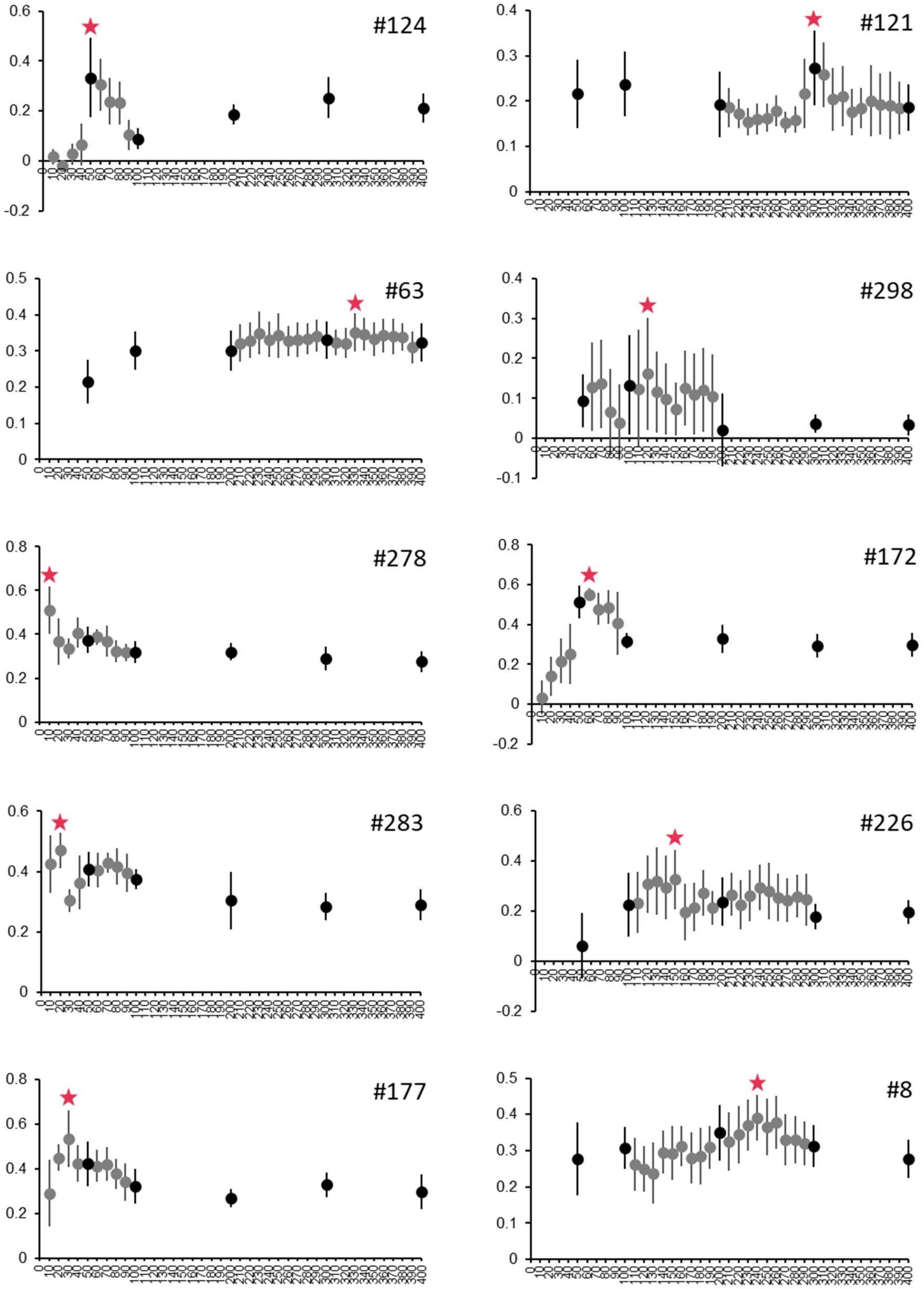

**Supplementary Fig. 1 (continued)**

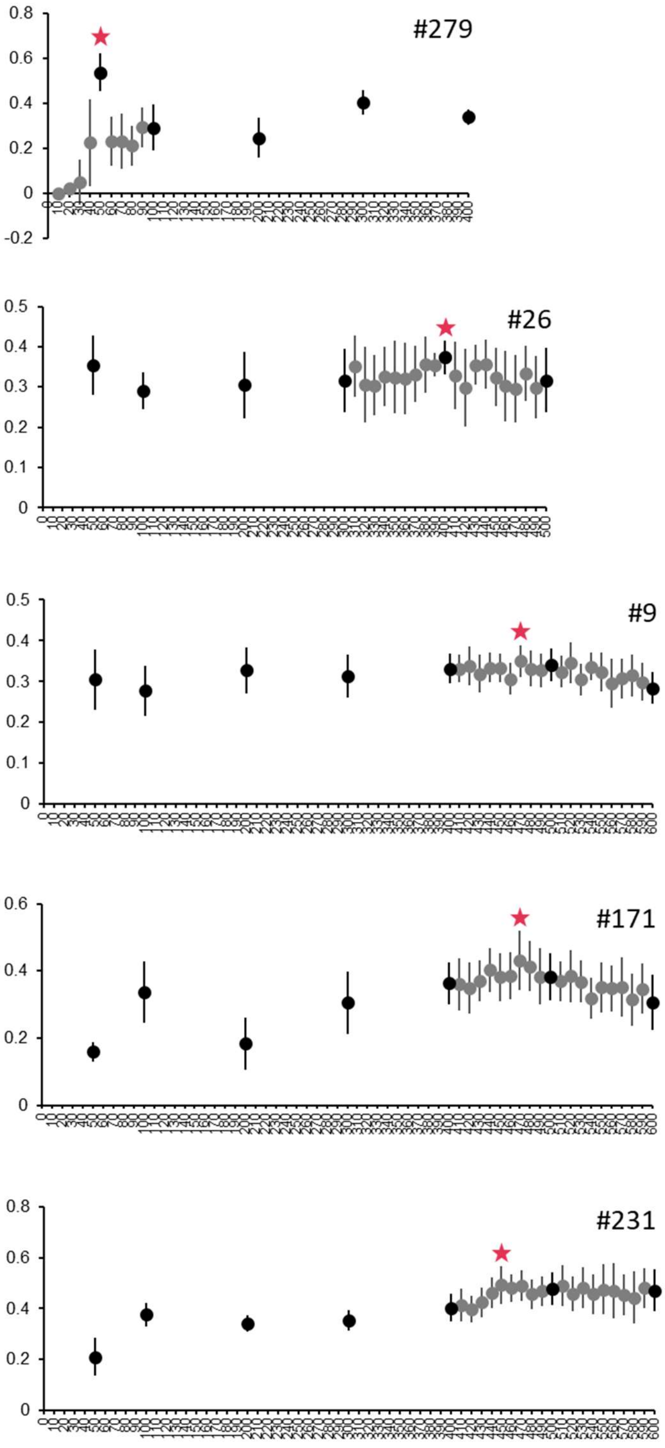

**Supplementary Fig. 1 (continued)**

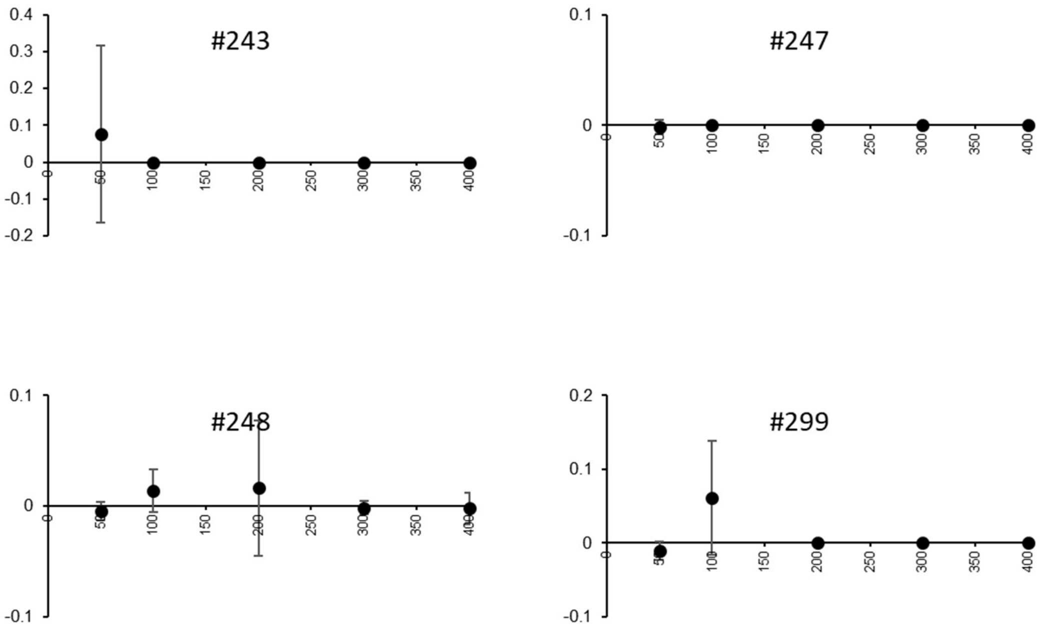

**Supplementary Fig. 1 (continued)** In multiple co-clustering using the top-ranked rs-FCs of these four MDD diagnostic biomarkers, the overall inter-dataset clustering stability was low, regardless of the number of rs-FCs.

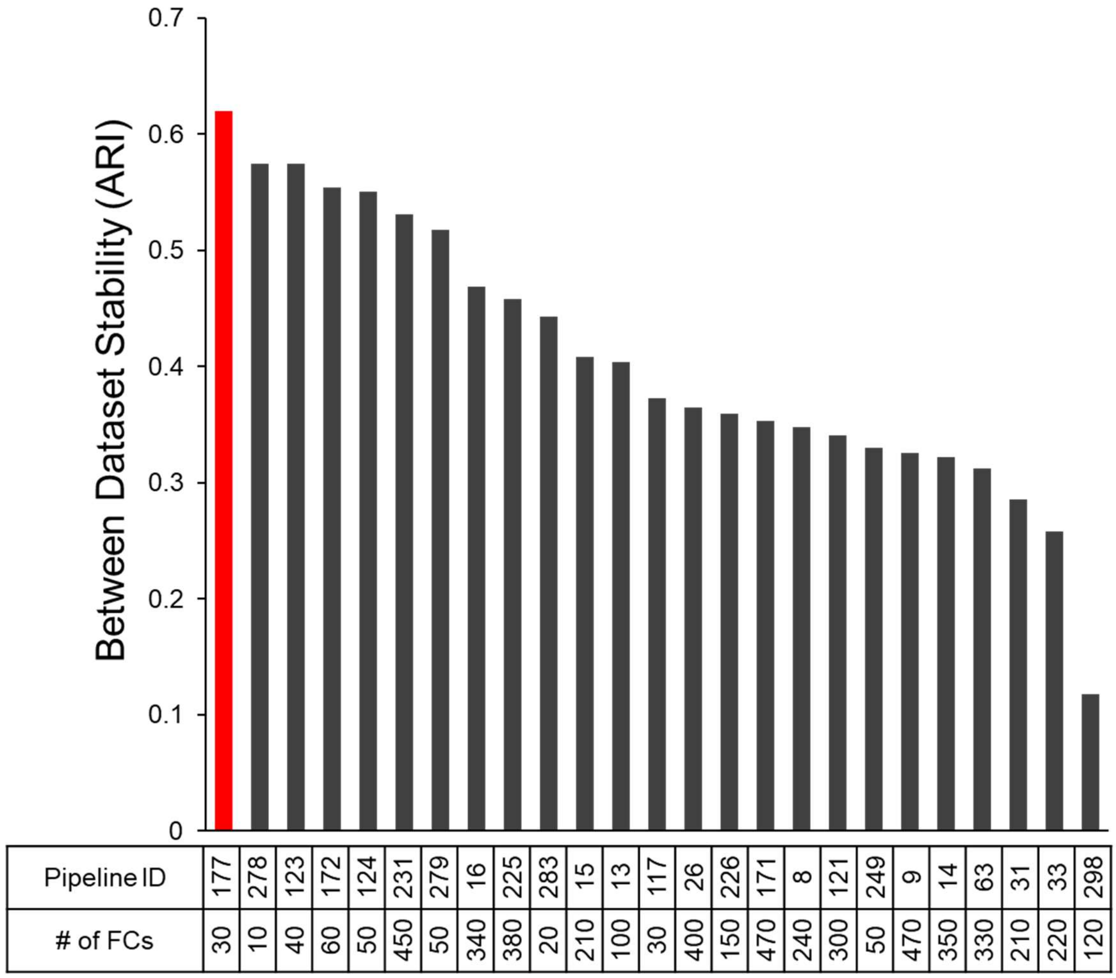

**Supplementary Fig. 2 The overall inter-dataset clustering stability obtained based on top-ranked rs-FCs of each diagnostic biomarker.** The horizontal axis shows the diagnostic biomarker construction pipeline IDs in Supplementary Table 1 and the numbers of top-ranked rs-FCs of MDD diagnostic biomarkers that showed the highest overall inter-dataset clustering stability (Supplementary Fig. 1).

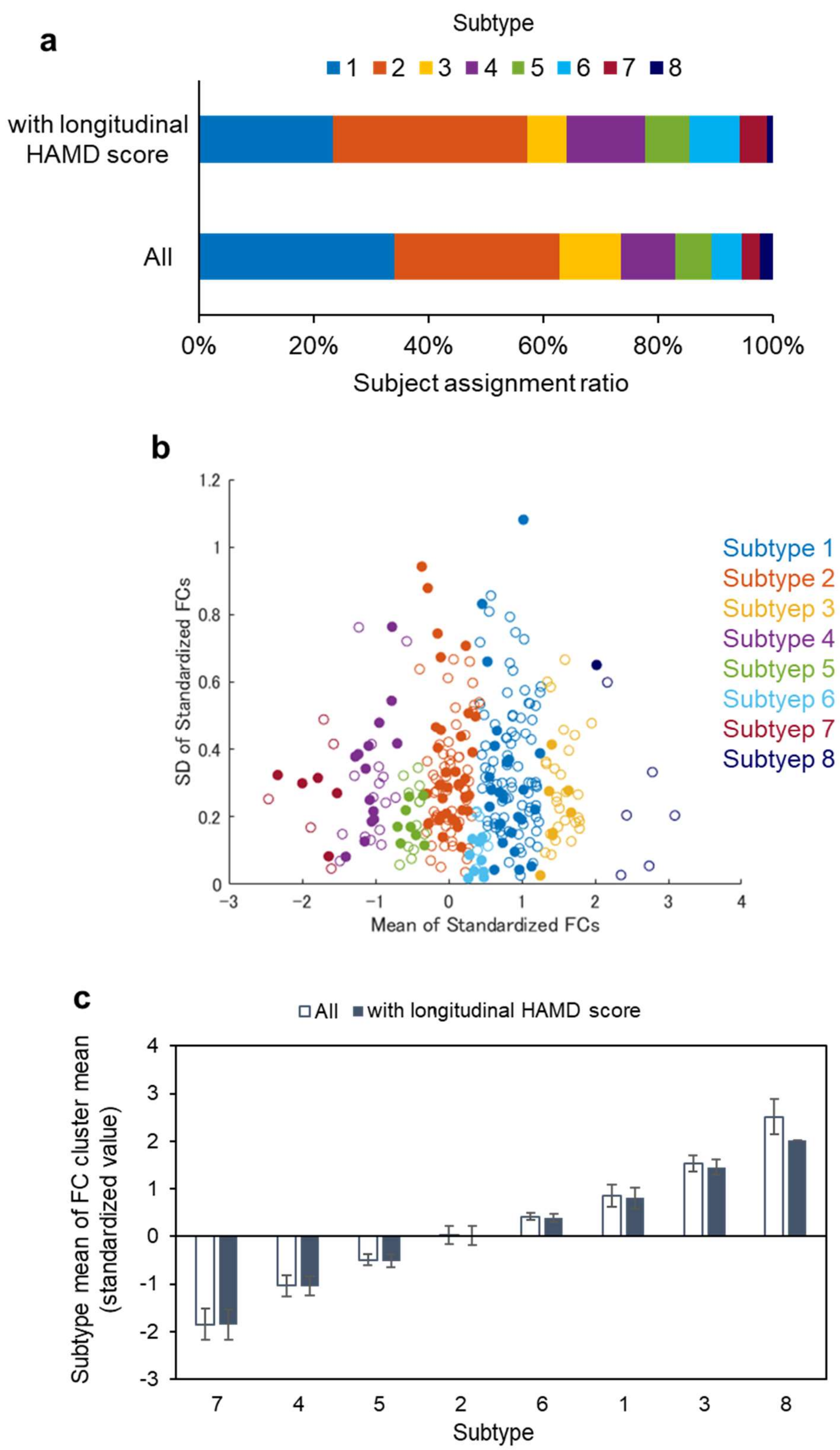

**Supplementary Fig. 3 Comparison of clustering structures and neurobiological characteristics of subtypes between all subjects and a subset of subjects with longitudinal clinical information.** a) Proportion of the number of people in each subtype. b) Visualization of the subject distribution of the three rs-FCs used in the stratification biomarkers. The horizontal axis shows the mean, and the vertical axis shows the standard deviation. The rs-FC values were standardized using subject data from each facility. Filled circle: Patient data accompanied by longitudinal clinical data. c) The averages of the three rs-FC averages for each subtype.

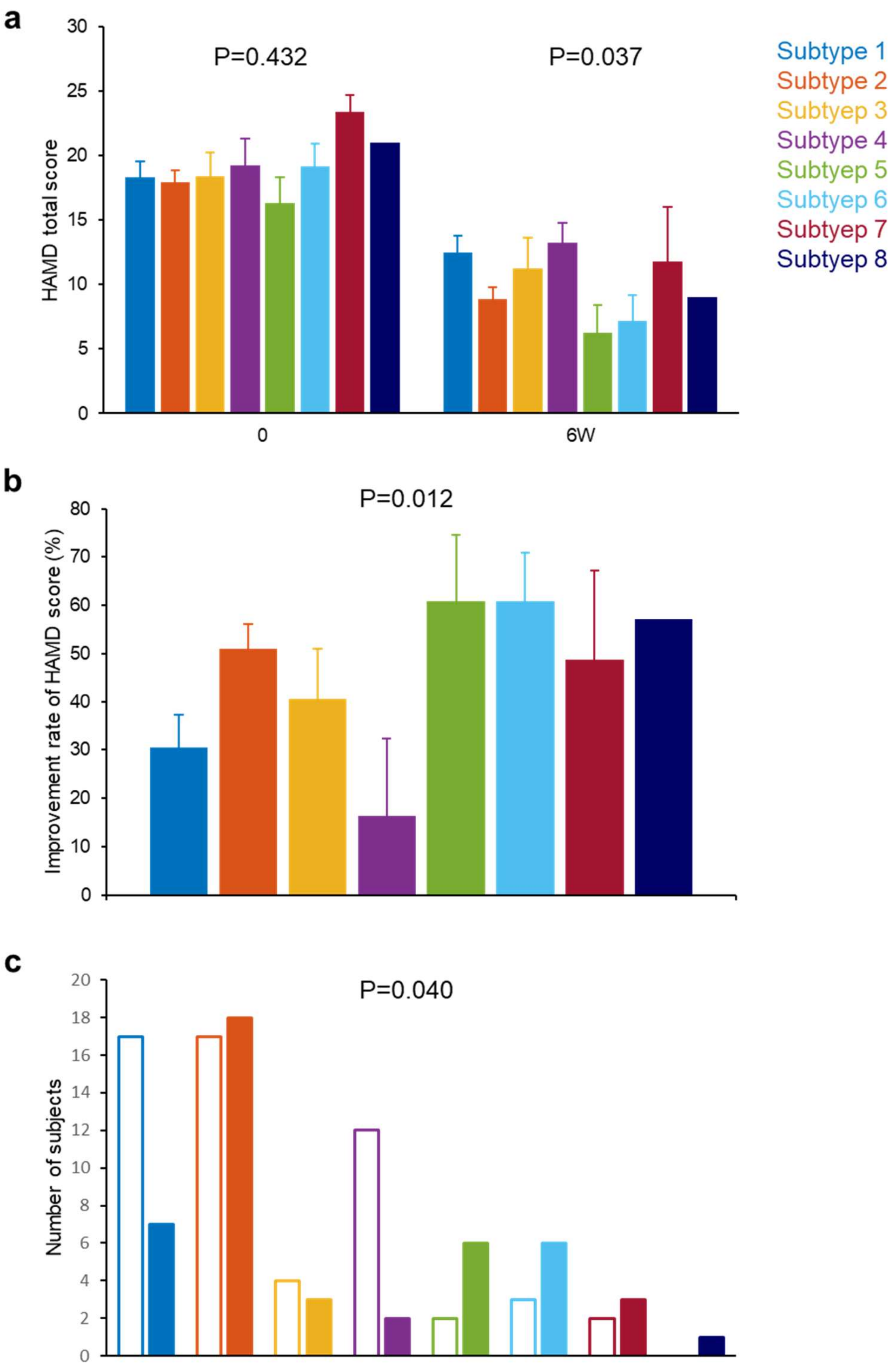

**Supplementary Fig. 4 Clinical characteristics analysis of all MDD subtypes.** a) The mean HAMD score of each subtype before and 6-8 weeks after the onset of treatment. P value by Kruskal-Wallis test. b) HAMD improvement rate of each subtype. HAMD improvement rate = (HAMD score before treatment - HAMD score 6-8 weeks after the onset of treatment) / HAMD score before treatment *100. P value by Kruskal-Wallis test. c) Response rate indicates percentage of subjects showing >50% reduction in HAMD score. P value by chi-square test.

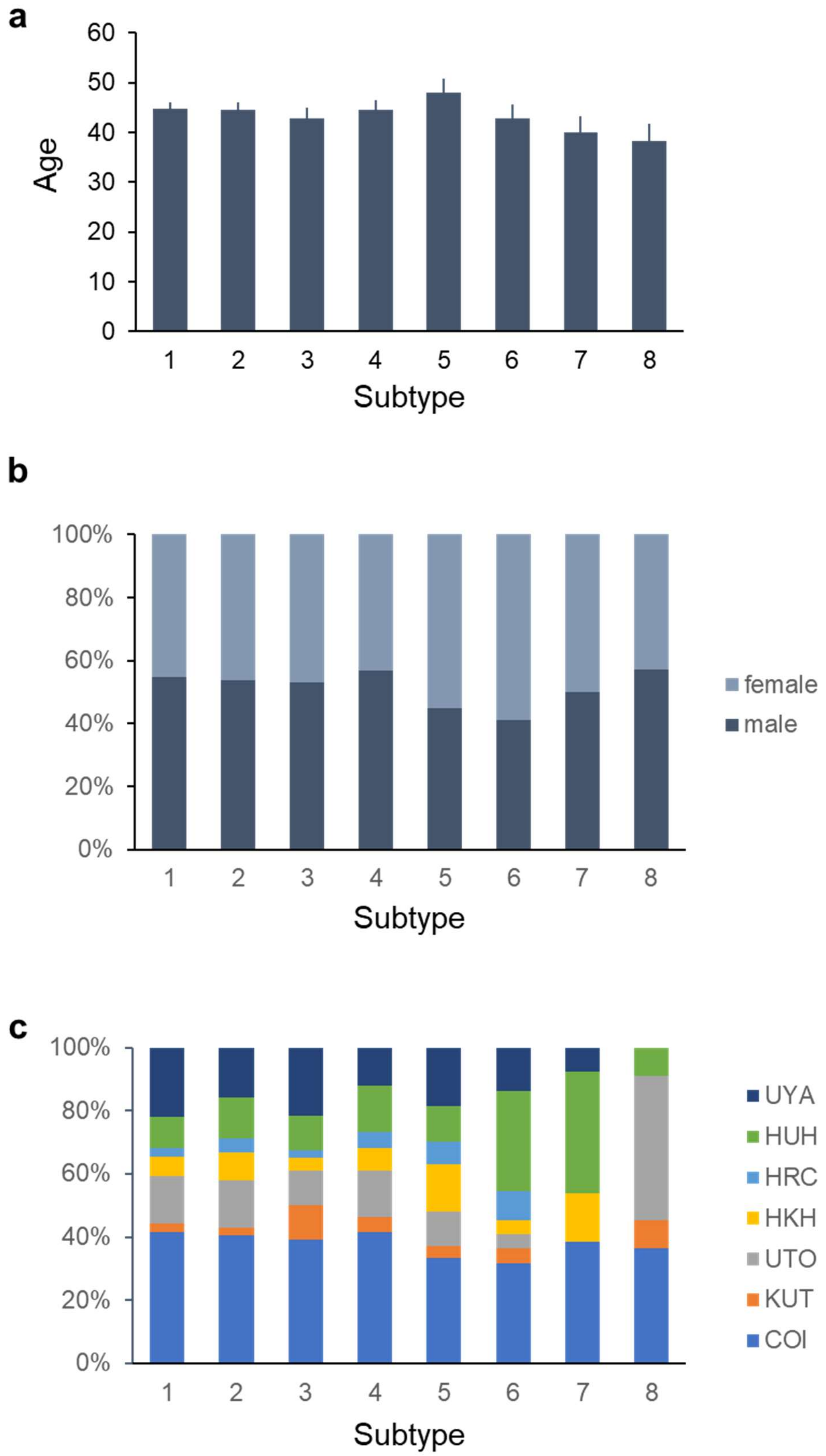

**Supplementary Fig. 5 Analysis of association with confounding factors.** a) Average age. b) Male-female ratio. c) Ratio of fMRI data acquisition facilities.

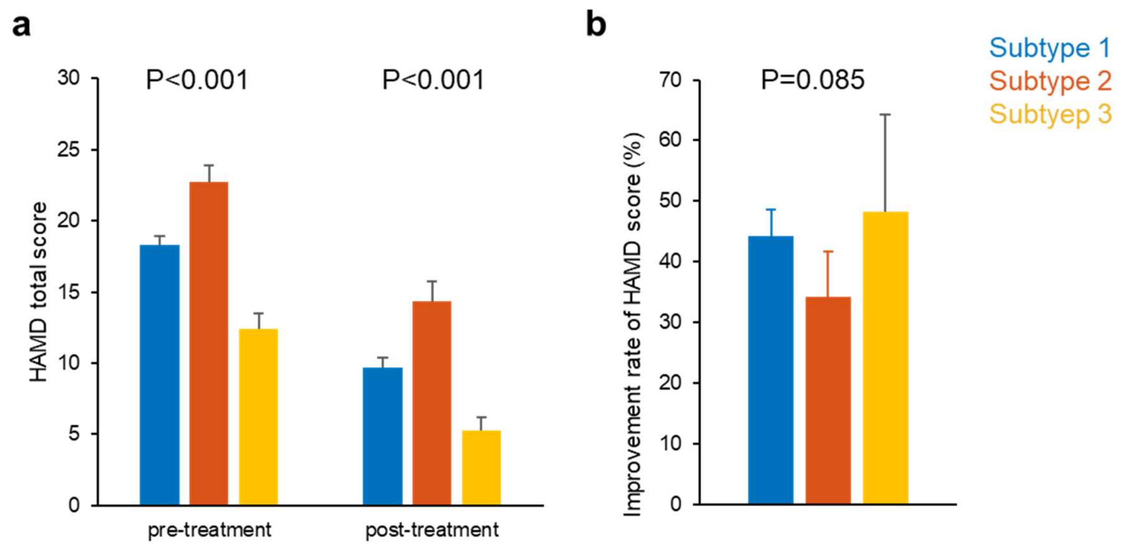

**Supplementary Fig. 6 Clinical characteristics of MDD subtypes stratified by clinical information.** a) The mean HAMD score of each subtype before and 6-8 weeks after the onset of treatment. b) HAMD improvement rate of each subtype. HAMD improvement rate = (HAMD score before treatment - HAMD score 6-8 weeks after the onset of treatment) / HAMD score before treatment *100. P value by Kruskal-Wallis test.
